# Large-scale population genomics of Malayan pangolins reveals deep diversification and a new species

**DOI:** 10.1101/2023.08.07.548787

**Authors:** Bo Li, Tianming Lan, Qing Wang, Minhui Shi, Ce Guo, Haimeng Li, Boyang Liu, Shanlin Liu, Kristen Finch, Shiqing Wang, Shangchen Yang, Liangyu Cui, Jun Li, Xilong Zhao, Jiangang Wang, Zhangwen Deng, Xinyu Wang, Yue Ma, Hyeon Jeong Kim, Samuel K Wasser, Kai Wang, Haorong Lu, Dongyi Yang, Jin Chen, Huabing Guo, Yan Yao, Hui Xie, Yiyi Wang, Jiale Fan, Wangsheng Li, Xiaotong Niu, Yinmeng Hou, Jieyao Yu, Jinyao Lu, Siyuan Li, Zhaowen Qiu, Wei Zhang, Suying Bai, Lei Han, Yuxin Wu, Xianchu Cai, Yunting Huang, Zhen Wang, Chen Wang, Jiahao Li, Yuze Jiang, Shasha Liu, Jiayi Wang, Li Li, Yan Hua, Huan Liu, Yanchun Xu

## Abstract

**Background:** Archipelagos and oceanic islands often present high percentage of endemism due to rapid speciation. The Malayan pangolin is a species distributing at both mainland (southern Yunnan, China) and oceanic islands via Malayan peninsula, which may result in deep differentiation among populations. In-depth investigation of population structure and genetic consequences for such species is of vital importance for their protection and conservation, practically for the critically endangered Malayan pangolin that is suffering from poaching, illegal trade, and habitat loss.

**Results:** Here we carried out a large-scale population genomic analysis for Malayan pangolins, and revealed three highly distinct genetic populations in this species, two of which are now being reported for the first time. Based on multiple lines of genomic and morphological evidence, we postulate the existence of a new pangolin species (*Manis*_1). Genetic diversity and recent inbreeding were both at a moderate level for both Malayan pangolins and *Manis*_1, but mainland Malayan pangolins presented relatively lower genetic diversity, higher inbreeding and fitness cost than island populations.

**Conclusions:** We found extremely deep and graded differentiation in Malayan pangolins, with two newly discovered genetic populations and a new pangolin species that is closely related to the Philippine pangolin than the typical Malayan pangolin, but a distant relative of the Indian pangolin. Anthropogenic factors did not significantly weaken the basis of genetic sustainability for Malayan pangolins, but mainland Malayan pangolins should be paid more attention for conservation due to higher genetic risks than island populations.

## Background

Comparing to continent, island biodiversity often presents high percentage of endemism^1^ due to rapid speciation^2–4^. For a species that widely distributes across mainland and islands may show deeper differentiation among island populations than mainland ones. Such geophylogenetic pattern requires special attention for conservation that local populations with unique evolutionary significance are risky to be overlooked given they were not properly identified. The Malayan pangolin (*Manis javanica*, Desmarest, 1822) is such a species distributing from mainland (southern Yunnan, China^5^) across Southeast Asian oceanic islands via Malayan pennisula^6^. It is reasonable to expect that in such large geographic range with complex ecological heterogeneity, deep differentiation would have happened.

Due to extensive habitat loss and consumption^7, 8^, this species declined towards the brink of collapse^9,10^, and was classified as Critically Endangered (CR) in the International Union for Conservation of Nature (IUCN) Red List^11^ in 2014, becoming of international conservation concern. The Convention on International Trade in Endangered Species of Wild Fauna and Flora (CITES) lists it in Appendix I^12^. Conservation is badly needed to simultaneously focus on poaching and illegal trade prevention, habitat and population restoration, and rescued animal rehabilitation^13, 14^. The effectiveness of conservation actions fundamentally depends on the background knowledge of fine-scale population structure^15^. However, studies on genetic differentiation of the Malayan pangolin is restricted to a few genome and genetic marker-based ones though researches on Asian pangolins has continued to advance, covering rehabilitation, captive breeding, anatomy and physiology, phylogenetics, evolution, ecology and trade^16^.

Hu et al. (2020a)^17^ analyzed 20.02 million genome-wide single nucleotide polymorphisms (SNPs) and sequence variations of mitochondrial Cytochrome c oxidase I (*COX1*) and Cytochrome b (*Cytb*) genes of 74 Malayan pangolins. They showed that Malayan pangolins could be divided into two main clades, i.e., MJA from the main continent and the Indochina peninsula and MJB from islands of Southeast Asia (exclusive to Java). The two main clades further differentiated into two and three subclades, respectively. However, the range of the Malayan pangolin holds very high ecological heterogeneity, where genetic differentiation of species is possible^18–20^. These regions have been recognized as a hotspot harboring some of the highest levels of biodiversity in the world^21^, potentially including additional pangolin species diversity yet to be described. For example, the Philippine pangolin (*M. culionensis*) used to be regarded as a subspecies of the Malayan pangolin, and was later defined as an independent species based on significant morphological and genetic differences resulting from long-term geographic separation^22–24^. In particular, Zhang et al. (2015)^25^ observed a unique monophyletic clade in *Manis*, and suspected that it was either a clade of the Philippine pangolin or possibly a new species that has not yet been discovered in the range of Malayan pangolin. Based on analysis of *COX1* and *Cytb* sequences of scale samples from Southeast Asia, Hu et al. (2020b)^26^ further proposed a potential fifth Asian pangolin species and estimated it diverged from Malayan pangolins 6.95 [4.64–9.85] million years ago (MYA) from Late Miocene to Early Pliocene. All these studies suggested that genetic divergence among populations of the Malayan pangolin might be deeper and more complex than previously recognized, even including undiscovered lineages and new species that could be supported with additional analyses of pangolin scales.

In this study, we re-sequenced 594 pangolin genomes, 10 of which were vouchered specimen of Malayan pangolin with known geographic origin and 584 of which were trafficked from Southeast Asia and confiscated by Chinese Customs over the last 20 years and have been identified as the Malayan pangolin based on morphological characteristics. By combining our data and genomic data from 73 previously published pangolin specimens^17^, we systematically analyzed population differentiation and fine-scale genetic structure by using the ever largest dataset and provided genomic and morphological evidence for the existence of a new species. Based on genomic analysis, we also assessed the potential genetic risk for the population. Our results provide novel implications for global conservation of the Malayan pangolin and new pangolin species.

## Results

### Genome resequencing and variants calling

We re-sequenced genomes from 594 pangolins, including 584 Malayan pangolins, one giant pangolin (*Smutsia gigantea*), three African tree pangolins (*Phataginus tricuspis*), four Indian pangolins (*M. crassicaudata*) and two Chinese pangolins (*M. pentadactyla*) (**Additional File 1: Table S1**). Thirty-nine samples with coverage below 80% were discarded from the whole genome analyses (**Additional File 1: Table S1**). The clean re-sequencing data of the remaining samples showed an average sequencing coverage of 91.23±0.74% and average sequencing depth of 16.65±3.74-fold. Sequencing data from each sample were mapped to the corresponding genomes, with acceptable mapping rate averaging 94.44±9.63%. A total of ∼95 million variants were detected, including 81,270,694 SNPs and 14,950,013 InDels. Additionally, we retrieved a consensus sequence of the whole mitochondrial genome for each pangolin individual.

### Deep differentiation in the Malayan pangolin and evidence for a new species

#### Phylogenetic analysis of Asian pangolins

To infer phylogenetic relationships of Asian pangolins, we constructed a Maximum Likelihood (ML) tree by combining our sequenced 525 genomes, the previously published 72 genomes in Hu et al. (2020a)^17^ and in Choo et al. (2016)^27^ (**Fig. 1a** and **Additional File 1: Tables S1–2)**. This phylogenetic relationship was also visualized using a simplified ML phylogenetic tree by randomly selecting 10 Chinese pangolins, 10 Malayan pangolins, 4 Indian pangolins, MJ-DCW-89, MJ-DCW-116, PA, and a dog as the outgroup (**Additional File 1: Fig. S1a**). The ML tree showed the phylogenetic relationships among Asian pangolins was *M. pentadactyla* + (*M. crassicaudata* + *M. javanica*). Interestingly, we found two samples, MJ-DCW-89 and MJ-DCW-116, forming a unique monophyletic group positioned outside Malayan pangolin with 100% bootstrap support. These results were also supported by principal component analysis (PCA) and admixture analysis (**Additional File 1: Fig. S1b–c**). Hereafter this clade is referred to as *Manis*_1 (*N* = 2). The remaining 596 Malayan pangolins (denoted as the typical Malayan pangolin) formed three distinct genetic clusters with deep divergence in the large-scale population. These three groups are denoted as MJ2 (*N* = 7), MJ3 (*N* = 22) and MJ4 (*N* = 567) (**Additional File 1: Table S1**). The clade MJ4 was further differentiated into 12 subclades (**Fig. 1d**). These results indicated deep diversification in Malayan pangolins.

**Fig. 1.**
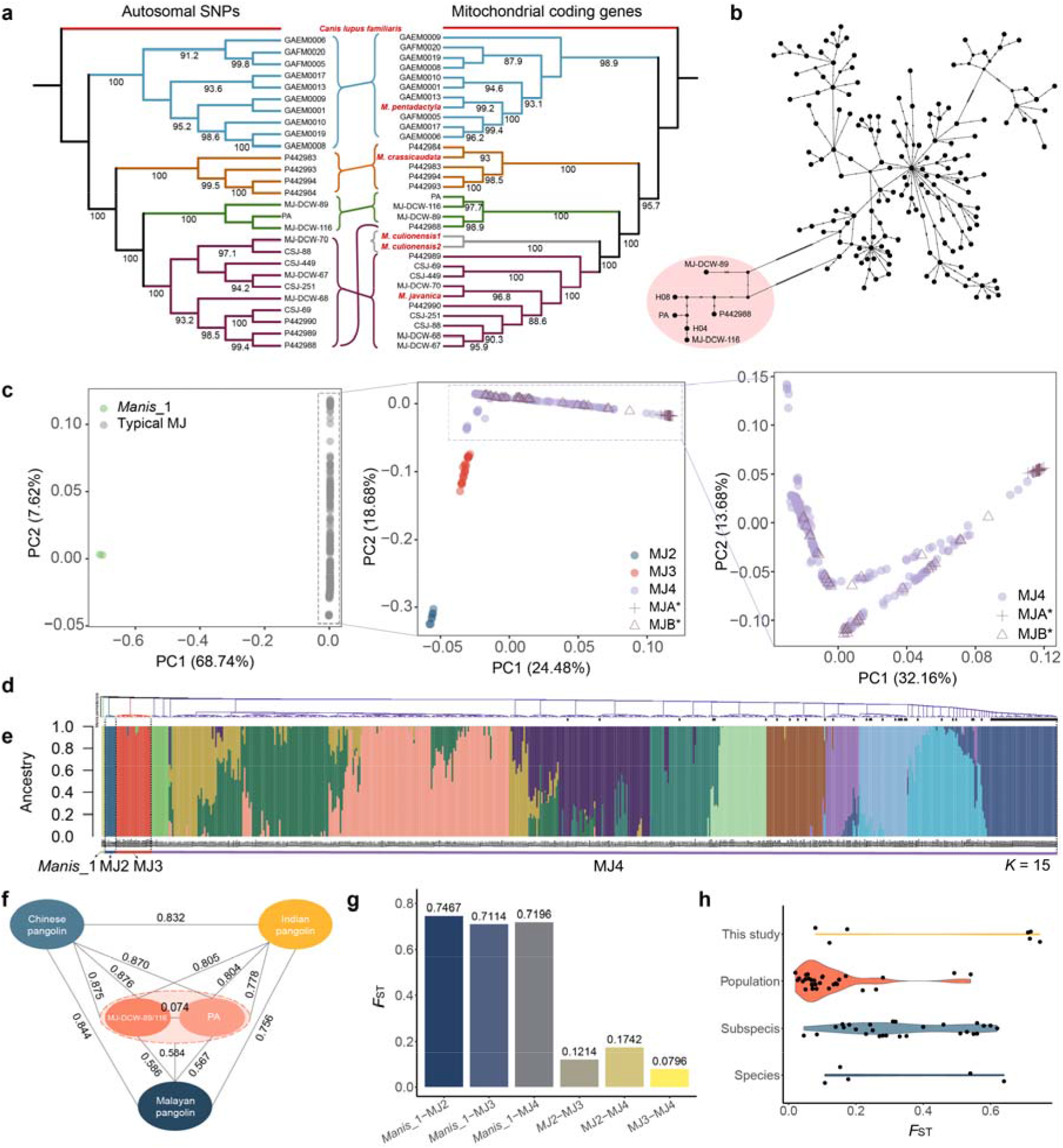
Genetic structure and differentiation of Malayan pangolins. **a** Maximum likelihood (ML) phylogenetic trees based on 13 mitochondrial coding genes (right) and autosomal SNPs (left). Incongruences between the two phylogenetic topologies are highlighted with colored lines. **b** Network analysis of *COX1* and *Cytb* gene by using Median-Joining method. The most diverged haplotype group containing MJ-DCW-89, MJ-DCW-116, PA, H04 and H08 was marked in pink. **c** PCA of all 598 pangolin genomes (left), PCA of 596 pangolins after the two *Manis*_1 samples (middle) were discarded, and PCA of MJ4 clade consisting of 567 Malayan pangolins (right). **d** ML phylogenetic tree based on the genome-wide SNPs with Chinese pangolin (*M. pentadactyla*) serving as an outgroup. **e,** Admixture analysis of all 598 pangolins (*K* = 15). **f** Pairwise *F*ST between each pair of Chinese pangolin, Indian pangolin, Malayan pangolin, PA and *Manis*_1. **g** Pairwise *F*_ST_ between each pair of four clades in this study. **h** Comparisons of pairwise *F*_ST_ in this study with selected species at population, subspecies, and species level.

We further constructed a ML tree using 13 concatenated mitochondrial gene sequences of 685 pangolins and 2 outgroup species to confirm this diversification (**Fig. 1a** and **Additional File 1: Fig. S2**). The phylogenetic relationships of these pangolin species were the same as previously reported by Gaubert et al. (2018)^24^, with African pangolins and Asian pangolins falling in two distinct clades. In the Asian clade, it showed similar phylogenetic structure inferred from whole genomes with the pattern *M. pentadactyla* + (*M. crassicaudata* + (*M. culionensis* + *M. javanica*)). As expected, *Manis*_1 formed a distinct monophyletic clade positioned between *M. crassicaudata* and the clade of *M. culionensis* + *M. javanica*. Malayan pangolins formed 17 distinct subclusters (**Additional File 1: Fig. S2**), demonstrating deep diversification. Furthermore, we reconstructed network figures (Network_660) using concatenated *COX1* and *Cytb* sequences. It showed the haplotypes from the *Manis_*1 clade (MJ-DCW-89 and MJ-DCW-116) clustered with the two haplotypes (H04: KT428139.1 and H08: KT428143.1; **Fig. 1b, Additional File 1: Fig. S3** and **Table S3**) that were suspected as a potential fifth Asian pangolin species in Zhang et al. (2015)^25^ and Hu et al. (2020b)^26^, and were position far away from main haplotype groups of Malayan pangolins. These results suggest that the clade *Manis_*1 represents a new pangolin species. Interestingly, the sample (P442988) that belonged to the *M. javanica* in the genome tree translocated to the clade *Manis*_1 in the mitochondrial genes tree (**Fig. 1b**).

#### Genetic structure

After removing 20 samples with high kinship score and 39 samples with low coverage sequences from our dataset (**Additional File 1: Tables S1** and **S4**), a total of 598 Malayan pangolin genomes containing 1,012,468 SNPs (after thinning) were used for population structure analysis. The principal component analysis (PCA) showed that the first principal component (PC1) separated *Manis_*1 samples from the rest 596 samples with 68.74% contribution rate (**Fig. 1c**). When the two representatives of *Manis_*1 were removed, the rest samples were clustered into three groups by the first and second components (PC1 = 24.48%, PC2 = 18.68%), corresponding to the MJ2, MJ3, and MJ4 identified in the phylogenetic tree (**Fig. 1d**). It is notable that MJ4 was highly complicated in genetic structure as demonstrated in the PCA and phylogenetic analyses. The clade MJ4 (**Fig. 1c**) contained 72 samples of the clade MJA and MJB defined in Hu et al. (2020a)^17^. However, the MJA and MJB were connected and formed an integrative meta-population when more samples were added (**Fig. 1c**).

Genetic structure was also supported by admixture analysis (**Additional File 1: Fig. S4**): the *Manis_*1 was firstly separated as an independent cluster when K ≥ 6, and the clade MJ2, MJ3 were all supported to be independent of MJ4 when K = 15 (**Fig. 1e**). F3 statistics further showed that F3 scores calculated for all 12 combinations of clusters were all positive, suggesting none of the four populations showed admixture with any other two populations of the same combination (**Additional File 1: Fig. S5**).

We calculated the *F*_ST_ among Asian pangolin species to test if the divergence between *Manis_*1 and other Malayan pangolins reached the species level (**Fig. 1f**). We found that the *F*_ST_ among Chinese, Indian, and typical Malayan pangolins ranged from 0.738 to 0.815. Taking this range as a reference of species-level divergence in Asian Pangolins, we found *F*_ST_ between *Manis_*1 and Chinese and Indian pangolins were 0.824 and 0.688, respectively. As we expected, the *F*_ST_ between the *Manis_*1 and three Malayan pangolin clades ranged from 0.711 to 0.747. This range is comparable with the *F*_ST_ range for recognized species and strongly supports our assertion that *Manis_*1 is a new pangolin species. Furthermore, we found the divergence between MJ2 and MJ3/MJ4 (*F*_ST(MJ2-MJ3)_ = 0.121, *F*_ST(MJ2-MJ4)_ = 0.174; **Fig. 1g**) has entered the range of subspecies level in some mammals like the Giant Panda^28^ and European rabbit^29^, though still less than those in most species such as Muskox^30^, African Green Monkey^31^, Bactrian Camel^32^, and Tiger^33, 34^ (**Fig. 1h** and **Additional File 1: Table S5**). Thus, differentiation of some local populations of typical Malayan pangolins appears to be approaching the subspecies level.

#### Morphometric variation of skulls

We collected 42 skulls from the sequenced pangolins, including MJ-DCW-116, the representative of *Manis*_1, three of the MJ2, seven of MJ3 and twenty-seven of MJ4 (**Additional File 1: Table S6**). The ratio of nasal bone length (NBL) to cranial length, which has been recommended as diagnostic characteristics to discriminate Malayan from Philippine pangolin, > 0.33 for Malayan and < 0.33 for Philippine^22^, ranged from 0.348 to 0.428 (averaged 0.386) for all our samples with only one marginally exceptional MJ4 sample (0.327). Virtually all of these skull measurements are accordant with Malayan pangolin. Ten traditional morphological indices were measured and compared with the data of previous studies^35^ (**Additional File 1: Table S7**). The PCA of these indices showed no significant differentiation of the four groups (**Additional File 1: Figs. S6–7** and **Table S8**).

By contrast, the skull of MJ-DCW-116 had significantly stronger and larger hamulus of pterygoid bone than the other 41 skulls compared to typical Malayan pangolin clades (**Fig. 2a**), and 13 of 75 landmarks on this skull deviate significantly from the average shape of typical Malayan pangolins at the anterior maxilla, orbital edge, the concavity at contact with the tympanic bulla, hypoglossus foramen, foramen magnum, the limit between the articular facet of the condyle and the foramen magnum, the intersection between inter-parietal and inter-frontal sutures, the intersection of frontal, parietal and squamosal, the intersection between interparietal suture and supraoccipital, the intersection between squamosal-parietal-exoccipital, and the condyle (**Fig. 2b** and **Additional File 1: Figs. S8–10**). When compared with the four Asian pangolin species, it also shows marked variation at anterior, sphenorbital fissure and the intersection of presphenoid, basisphenoid, and palatine (**Fig. 2c** and **Additional File 1: Fig. S11**).

**Fig. 2.**
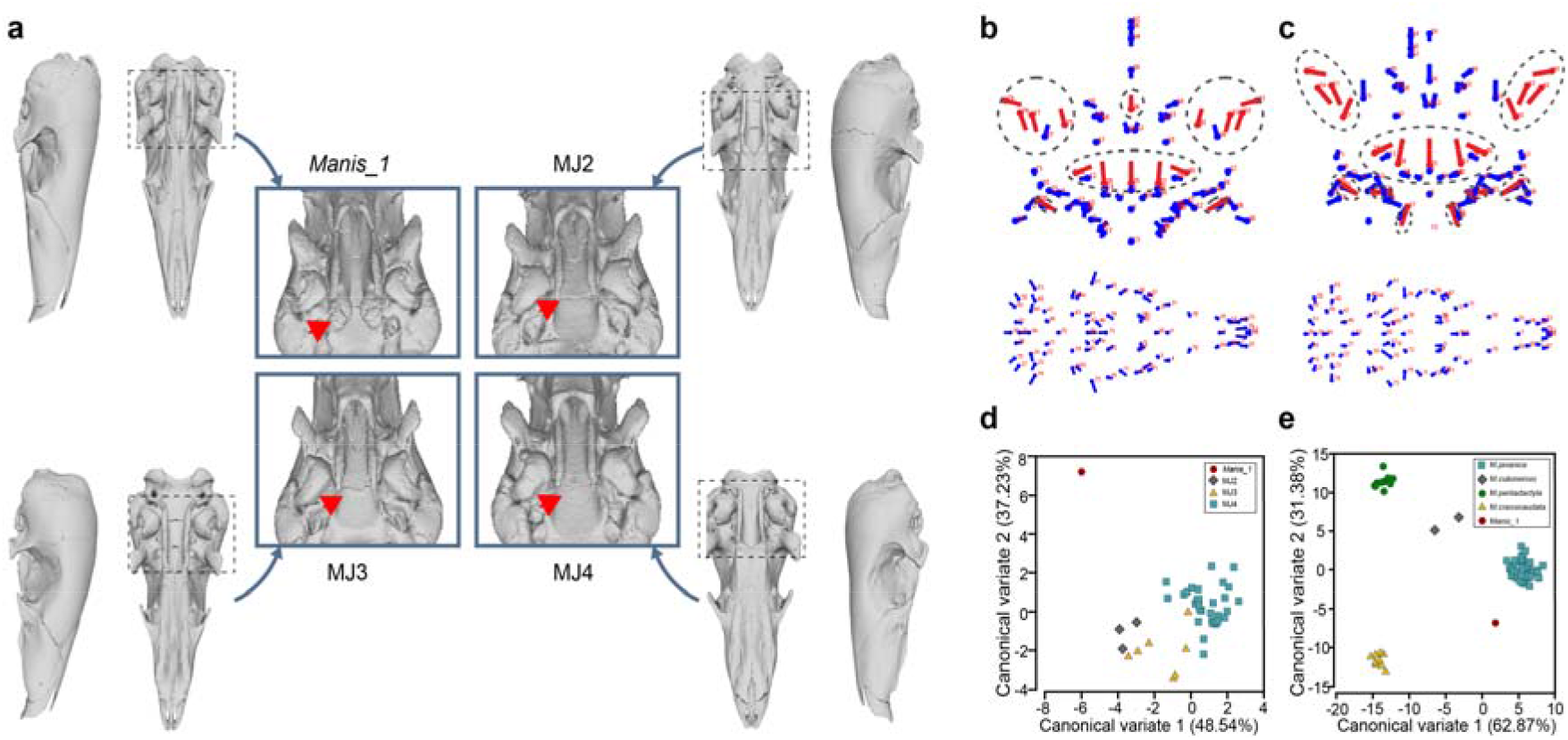
Morphology and morphometric analysis of pangolin skulls. **a** The skull of *Manis*_1 compared to skulls from Malayan pangolin subpopulations; The region that *Manis*_1 differ from typical Malayan pangolins is indicated by red arrows. **b** The upper figure shows the shape changes at landmarks on the *Manis*_1 skull (red lollipops in dashed circles) comparing to typical Malayan pangolins (blue lollipops). The lower figure shows the overall shape variation of typical Malayan pangolin skulls. **c** The upper figure shows the shape changes at landmarks on the *Manis*_1 skull (red lollipops in dashed circles) comparing to four species of Asian pangolin including the typical Malayan pangolin. The lower figure shows overall shape variation in the four Asian pangolin species including the typical Malayan pangolin. **d** Canonical variate scatter plot (CV1 and CV2) showing separation between the skull of *Manis*_1 and typical Malayan pangolins. **e** Canonical variate scatter plot showing the separation of skulls of *Manis*_1 and four species of Asian pangolins including the typical Malayan pangolin along the first two canonical variate (CV1 and CV2) axes.

The scatterplot of Canonical Variate Analysis (CVA) shows that the typical Malayan pangolin clades clustered in a large group and separated from *Manis*_1 represented by MJ-DCW-116 in morphological space (**Fig. 2d** and **Additional File 1: Fig. S12**). The first two canonical variates accounted for 85.77% total variation (CV1 = 48.54%, CV2 = 37.23%) (**Additional File 1: Table S9**). CV1 revealed significant shape variation at landmarks in the anterior of the ventral maxilla, the frontal-parietal-squamosal intersection, the squamosal-parietal-exoccipital junction and the supraoccipital portion of the skull roof. CV2 showed the shape variation at the posterior end of the premaxillary on the midline, the maxillary projection into the palatine, the optic foramen (**Additional File 1: Fig. S13**). For the four Asian pangolin species, the first two canonical variates accounted for 94.25% total variation (CV1L=L62.87%, CV2L=L31.38%) (**Additional File 1: Table S10)**, and all species, including *Manis*_1 could be distinctly separated from each other on the scatter plot (**Fig. 2e**). The inter-specific shape variation mainly occurred at the sphenorbital fissure and the anterior-most part of the zygomatic process of the squamosal as revealed by CV1, and the anterior of the ventral maxilla and the premaxillary on the midline as revealed by CV2 (**Additional File 1: Fig. S14**).

The Mahalanobis distances among typical Malayan pangolin clades MJ2–MJ4 ranged from 3.69 to 5.25 with a mean 4.45. The distances of *Manis*_1 to these clades varied from 9.60 to 10.27 and averaged 9.92 (**Additional File 1: Table S11**). Similarly, the Procrustes distances of *Manis*_1 to these clades varied from 0.076 to 0.079 and averaged 0.077, significantly greater than those among clades MJ2–MJ4 that varied from 0.025 to 0.043 with a mean 0.036 (**Additional File 1: Table S12**). For the four Asian pangolin species, the inter-specific Mahalanobis and Procrustes distances ranged from 16.92 to 22.98 (20.95±3.00) and from 0.068 to 0.1008 (0.091±0.013), respectively (**Additional File 1: Tables S13–14**). The two distances of *Manis*_1 to the four species were 15.79 to 27.35 with a mean 21.99±4.77 for Mahalanobis distances (**Additional File 1 Table S13**) and 0.076 to 0.145 with a mean 0.117±0.030 for Procrustes distances (**Additional File 1: Table S14**). Both fell in or even slightly exceeded the inter-specific ranges.

#### Inference of geographic origin of main clades

Here we attempt to infer the geographic origin of these confiscated pangolins by comparison to samples of known origins (**Fig. 3**). PCA analysis on nuclear genomes (**Fig. 3a**) showed that two Malaysian samples from northeast part of Borneo were assigned to MJ2, and one Malaysian sample from northwestern Borneo was assigned to MJ3. These suggest that the typical Malayan pangolin on Borneo differentiated into two regional populations corresponding to the clade MJ2 in the northeastern part and the MJ3 in northwestern part (**Fig. 3b**). Twelve known samples were also assigned to MJ4 including 3 from China, one from Myanmar, one from Cambodia, four from Singapore, one from Malaysia, and two from Indonesia (**Fig. 3a**). This suggested the Clade MJ4 could be a meta-population consisting of continent-peninsula-island (Sumatran) regional populations (**Fig. 3b**). None of our known samples were assigned to *Manis*_1 based on nuclear genomic analysis.

**Fig. 3.**
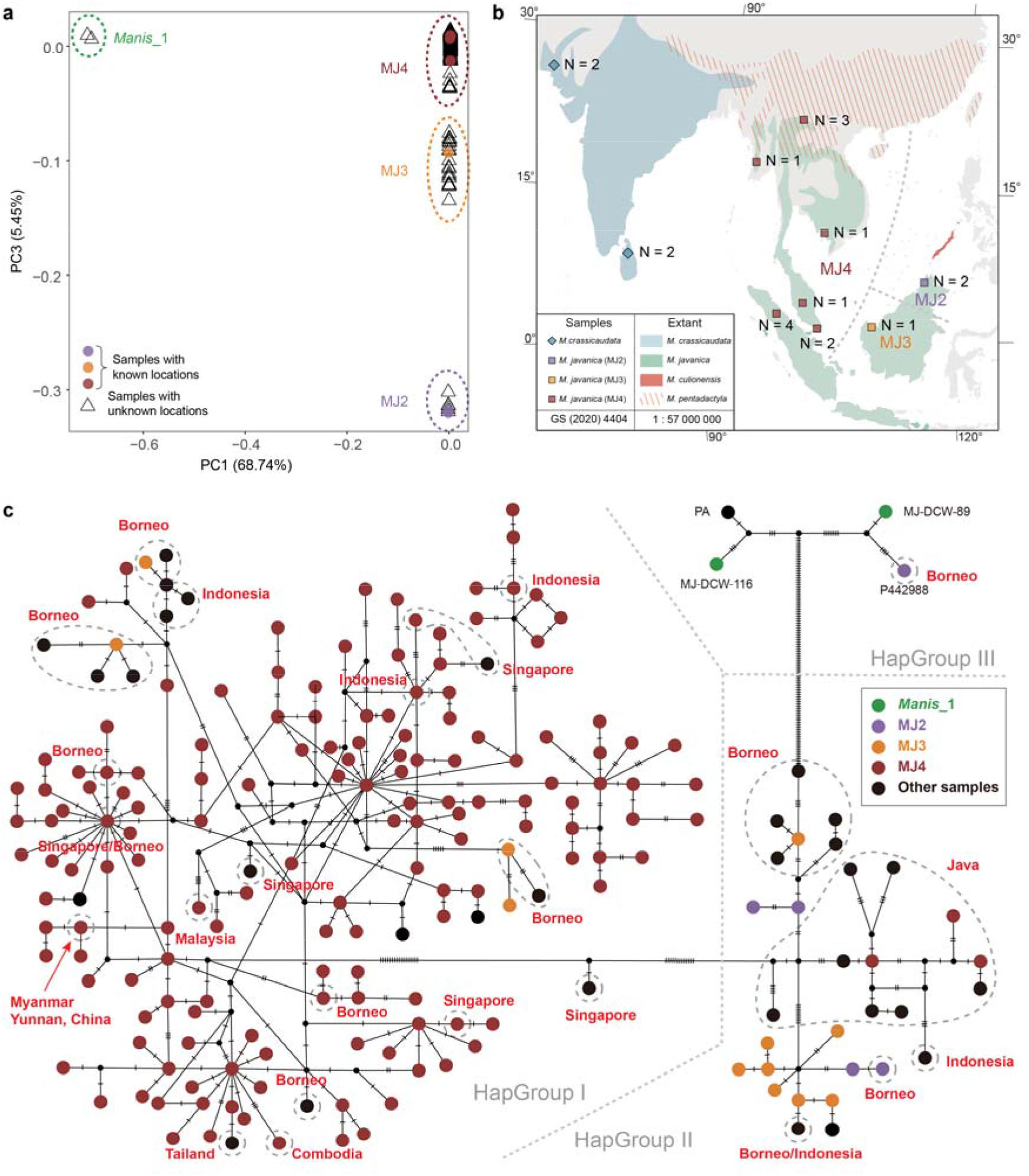
Geographic origin inference using principal component analysis of nuclear genomes and mitochondrial haplotype network. **a** PCA plot of samples based on nuclear genome. Samples with known sampling sites were marked as colored solid squares. **b** The geographic range of the typical Malayan pangolin showing the samples with known sampling sites. **c** Network of 205 mitochondrial haplotypes of typical Malayan pangolins and *Manis_*1. Haplotypes in gray dashed circles are the published haplotypes of Malayan pangolins with known sampling sites (*N* = 60). The three typical Malayan populations and new species defined in this study are shown in different colors.

We then reconstructed a haplotype network with concatenated *COX1* and *Cytb* sequences including previously published homologous sequences of 64 Malayan pangolins with known geographic locations (**Fig. 3c, Additional File 1: Fig. S15** and **Tables S15–16**). Haplotypes generally fell into three main groups: HapGroup I consisted of haplotypes from north Borneo, Singapore, Sumatra, Malaysia peninsular, and northward to Thailand and Myanmar, covering MJ4 and a part of MJ3; HapGroup II consisted of haplotypes from Java, south Borneo, Indonesia, and south Sumatra, covering MJ2 and a part of MJ3. HapGroup III included two *Manis*_1 haplotypes (MJ-DCW-89 and MJ-DCW-116) and a northeast Borneo haplotype from Malaysia (P442988). Such pattern firstly supports the chromosomal genomic inference that the typical Malayan pangolin across Southeast Asia exhibited significant genetic differentiation but did not reveal the proposed pattern of continental and island populations^17^, and secondly suggests the new species *Manis_*1 might be from regions around northeast Borneo.

### Population history of different clades

#### Gene flow among clades

Based on the phylogenetic relationship inferred using genome-wide SNPs and mitochondrial genes, we further investigated the signals of post-divergence gene flow between *Manis_*1 and the three typical Malayan clades through D-statistics with the Chinese pangolin as an outgroup (MP). We found that *Manis_*1 shared more alleles with MJ2 than MJ3/MJ4 (|Z-score| > 3), suggesting gene flow between *Manis_*1 and MJ2. We detected no significant gene flow between *Manis_*1 and MJ3/4 (|Z-score| < 3) (**Fig. 4a**). Identity-by-decent (IBD) analysis showed no shared segments greater than 1 Mb between *Manis_*1 and MJ2–4, and between MJ2 and MJ3 (**Fig. 4b**). For medium-size segments (between 100 kb and 1 Mb in length), we identified seven sections ranging from 0.33 to 0.40 Mb shared between MJ2 and MJ4, and 93 sections ranging from 0.30 to 0.89 Mb shared between MJ3 and MJ4.

**Fig. 4.**
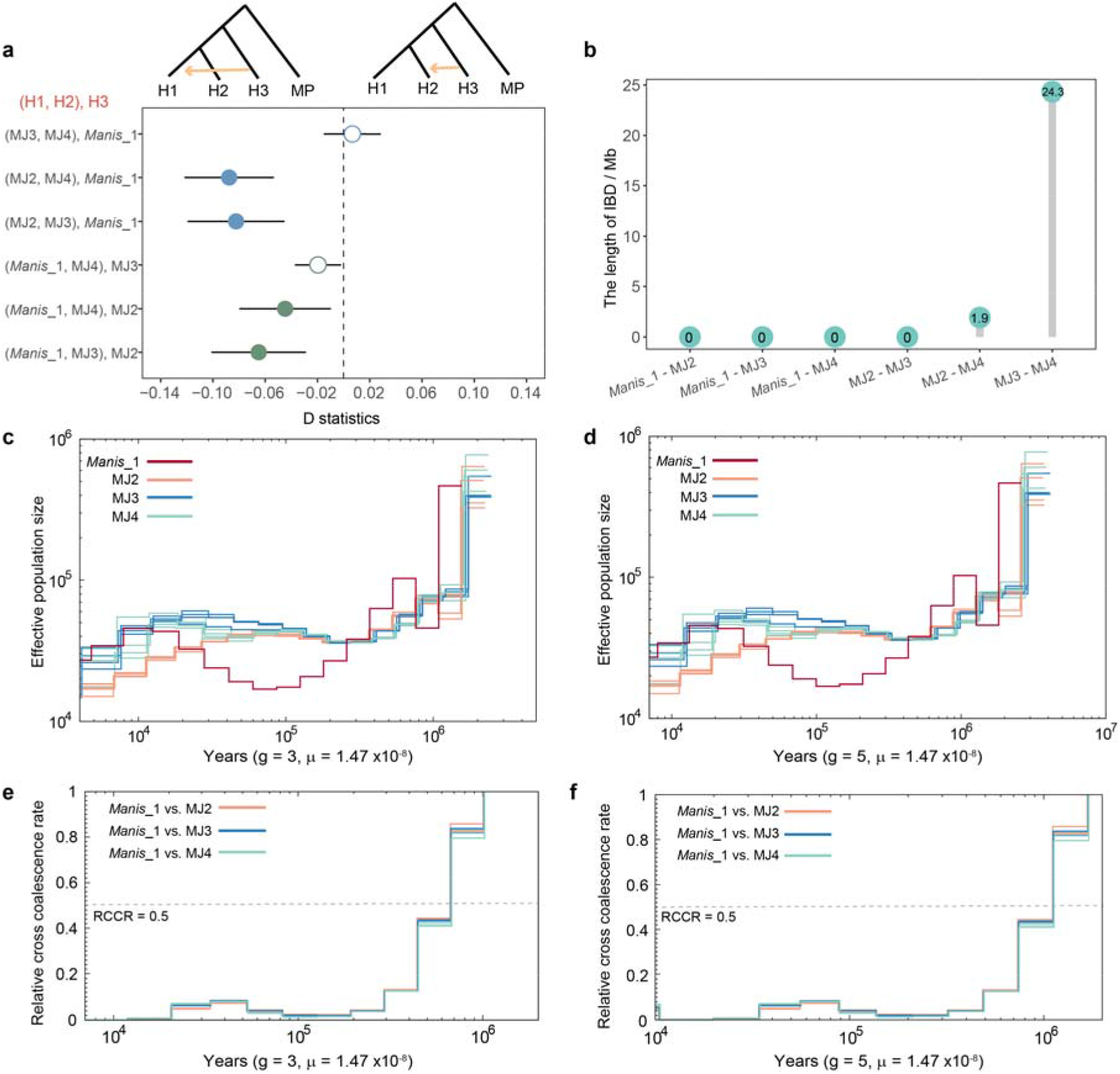
Gene flow and demographic history of populations. **a** D-statistics analysis with the Chinese pangolin as an outgroup. H1, H2 and H3 represents each population of MJ2–MJ4 and *Manis*_1. Each solid circle represents the mean absolute Z-score more than 3. Each hollow circle represents the mean absolute Z-score less than 3. The dashed vertical line denotes D = 0. **b** The length of IBD segments shared between each pair of *Manis*_1 and three typical Malayan pangolin clades. **c, d** Inferred effective population sizes over time for *Manis*_1 and each of three typical Malayan pangolin clades using MSMC2 with generation length setting as 3 years (**c**) and 5 years (**d**). **e, f** Inferred divergence time of pangolin clades using MSMC2 with generation length setting as 3 years (**e**) and 5 years (**f**).

#### Demography of different clades

By using MSMC2 analysis, the demographic history of the typical Malayan pangolin and the new species *Manis_*1 was traced back to approximately 3–2 million years ago (Mya) during which period they experienced distinct demographic histories (**Fig. 4c–d**). The three typical Malayan pangolin clades experienced similar demographic trajectories with three phases: 1) a sharp population decline from 3.0–2.0 Mya to 0.6–0.4 Mya; 2) followed by a gradual expansion and a peak approximately 40–20 thousand years ago (kya); and 3) a final decline to the present-day. However, the clade MJ2 in comparison did not restore as much and continued to decline approximately 45–27 kya potentially leading to its divergence from the clade MJ3 and MJ4 (**Additional File 1: Fig. S16**).

In contrast, the new species underwent a different demographic history from the typical Malayan pangolin with two bottlenecks and two large expansions. The population declined sharply to the first bottleneck as the typical Malayan pangolin did but more recently 1.8–0.9 Mya (**Fig. 4c–d**). It appears to have rebounded rapidly, but soon slid into the second bottleneck approximately 150–90 kya. Afterward, the population restored to the similar level as the typical Malayan pangolins until 20–10 kya followed by a recent synchronized decline. The estimated divergence time of *Manis*_1 from the typical Malayan pangolin was about 1.2–0.7 Mya when RCCR = 0.5 (**Fig. 4e–f**).

### Evaluation of evolutionary potential of typical Malayan pangolins and the new species

#### Genetic diversity and inbreeding

We used nucleotide diversity (π) and average heterozygosity (*He*) to estimate genetic diversity for the new species and the typical Malayan pangolins. The π value for all typical Malayan pangolins was 0.00234±0.00139 (**Additional File 1: Fig. S17a)**, 0.00219±0.00150 for MJ2, 0.00244±0.00152 for MJ3 and 0.00231±0.00138 for MJ4. This value was 0.00128±0.00142 for *Manis*_1, lower than those for typical Malayan pangolins. The *He* for all typical Malayan pangolins was 0.00183±0.00013 (**Additional File 1: Fig. S17b**), 0.00187±0.00008 for MJ2, 0.00210±0.00005 for MJ3 and 0.00181±0.00041 for MJ4. Similar to π value, the new species possessed much lower *He* (0.00134±0.00000). However, the genome-wide mean *He* of *Manis*_1 and typical Malayan pangolins were at medium or high level among endangered species (**Fig. 5a** and **Additional File 1: Table S17**). The genetic diversity of the new pangolin species was comparable to the critically Endangered (CR) Chinese pangolin (*He* = 0.00182), vulnerable (VN) Giant panda (*Ailuropoda melanoleuca*), and western lowland gorilla (*Gorilla gorilla gorilla*, CR) etc., and higher than the Iberian lynx (*Lynx pardinus*, EN), Island fox (*Urocyon littoralis*, CR) and African cheetah (*Acinonyx jubatus*, VN) *etc*. (**Fig. 5a** and **Additional File 1: Table S17**). It was noteworthy that a group of samples of MJ4 had lower *He* compared with all other pangolins, at the similar level of the tiger (**Fig. 5a**). They were from the mainland Asia, including China and Myanmar.

**Fig.5.**
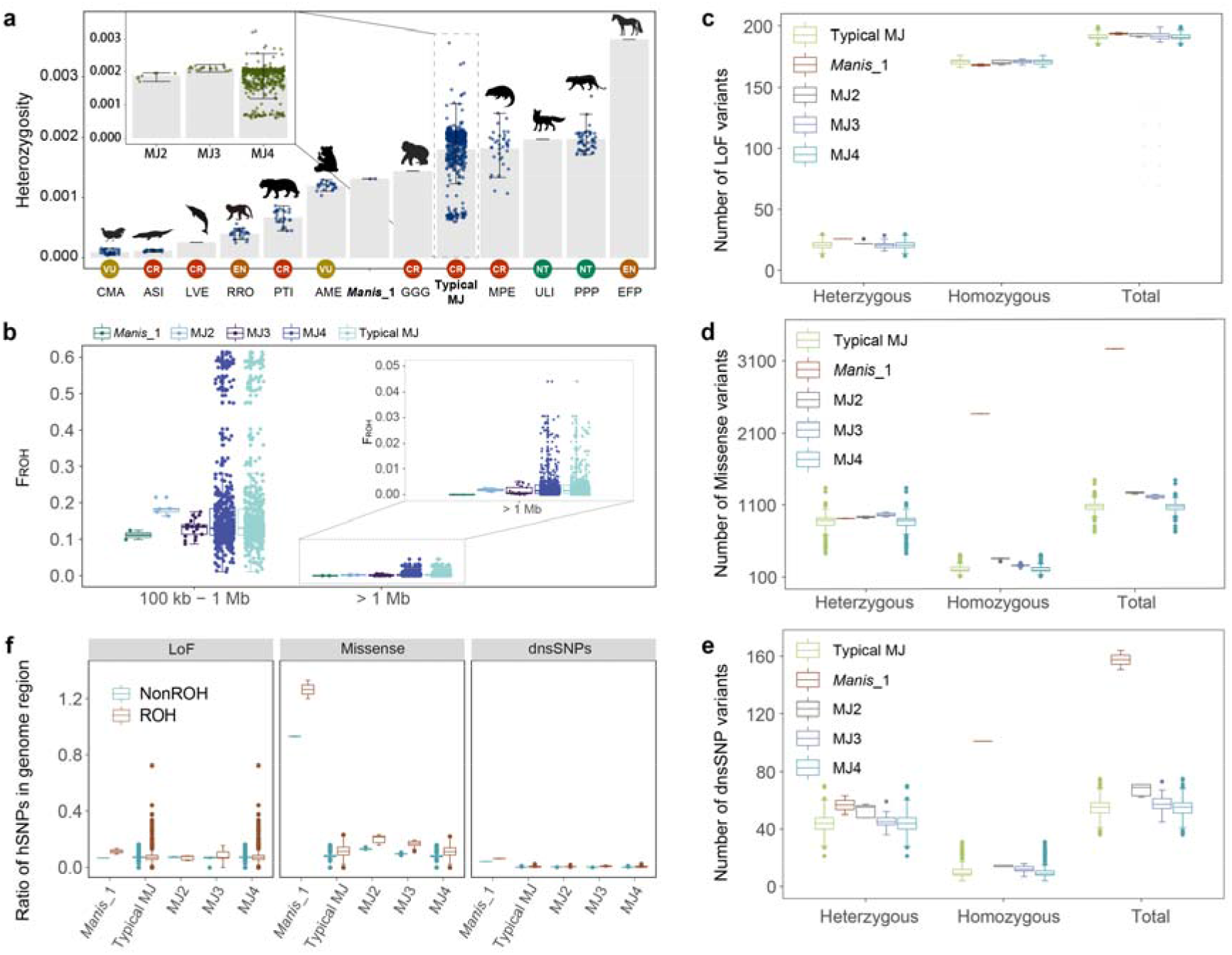
Genetic diversity, ROH, and mutational loads in the three typical Malayan pangolin clades and *Manis*_1. **a** Comparison of genome-wide heterozygosity of the three Malayan pangolin clades and the new species *Manis*_1 with reference to other endangered species. Colored dots represent individuals and whiskers represent the 95% confidence interval of the genome-wide heterozygosity in each species. X-axis abbreviations are as follows: CMA: Brown eared pheasant (*Crossoptilon mantchuricum*), ASI: Chinese alligator (*Alligator sinensis*), LVE: Baiji (*Lipotes vexillifer*), RRO: Golden snub-nosed monkey (*Rhinopithecus roxellana*), PTI: South China tiger (*Panthera tigris amoyensis*) and Siberian tiger (*P. t. altaica*), AME: Giant panda (*Ailuropoda melanoleuca*), GGG: Western lowland gorilla (*Gorilla gorilla gorilla*), MPE: Chinese pangolin (*Manis pentadactyla*), ULI: Island fox (*Urocyon littoralis*), PPP: African leopard (*Panthera pardus pardus*), EFP: Przewalski’s horse (*Equus ferus przewalskii*). **b** Fraction of ROH (*F*_ROH_) with different length categories (100 kb < ROH < 1 Mb and ROH > 1 Mb) in the genomes of *Manis*_1 and typical Malayan pangolins. *Manis*_1: the new pangolin species, MJ2–4: three typical Malayan pangolin clades, and Typical MJ: the total typical Malayan pangolins. **c, d** Comparison of the number of heterozygous, homozygous, and total Loss-of-Function (LoF, **c**) SNPs, missense SNPs (**d**) and deleterious nonsynonymous SNPs (dnsSNPs, **e**). **f** Comparison of the ratio of homozygous LoF, missense and dnsSNPs variants (hSNPs) in the ROH, and non-ROH regions of each individual’s genome.

We also tested the inbreeding level in different clades by screening runs of homozygosity (ROHs) across the whole genome. The medium-sized ROHs (100 kb to 1 Mb) of the typical Malayan pangolin averaged 419.6±302.4Mb, and the long-sized ROHs (> 1 Mb) averaged 8.3 Mb±13.4 Mb. The mean size of medium-and long-sized ROHs in clade MJ2 were 447.1±42.7 Mb and 4.6±1.4 Mb, respectively, and the two sized ROHs in clade MJ3 and MJ4 were 319.7±57.2 Mb, 423.1±308.5 Mb and 3.8±4.1 Mb, 8.6±13.7 Mb, respectively. For the new species *Manis*_1, we did not detect long-size ROHs from the two representatives, but the medium-size ROHs averaged 272.4±31.6 Mb (**Additional File 1: Fig. S18**). The mean medium-size *F*_ROH_ of *Manis*_1 was 11.17±1.30%. This *F*_ROH_ was 17.18±12.40% for typical Malayan pangolins, 18.33±1.75% for clade MJ2, and 13.11±2.35%, and 17.35±12.65% for clade MJ3 and MJ4. The mean long-sized *F*_ROH_ of the new species was 0, and of the typical pangolin was 0.34±0.55%, of which *F*_ROH_ was 0.19±0.06% for clade MJ2, and 0.15±0.17%, 0.35±0.56% for clade MJ3 and MJ4 (**Fig. 5b**). The only group of samples with a very high *F*_ROH_ was from mainland Asia, including China and Myanmar (**Fig. 5b**). In general, the inbreeding level in *Manis*_1 and typical Malayan pangolins were low, with the only exception of mainland Malayan pangolin (Myanmar and Yunnan, China, **Fig. 5b**).

#### Mutational load

We screened mutational loads by counting loss-of-function mutations (LoFs), missense mutations, and deleterious nonsynonymous SNPs (dnsSNPs) on each individual genome. To reduce the calculation bias introduced by the reference genome, we used the cat genome as a reference to detect derived mutational loads. We found very similar number of LoFs in each of the three typical Malayan clades and the new species (MJ2: 193±1; MJ3: 192±3; MJ4: 192±3; *Manis*_1: 194±1) (**Fig. 5c**). For missense mutation and dnsSNPs, however, *Manis*_1 presented an excess of mutational loads compared to typical Malayan pangolins (**Fig. 5d–e)**. The number of missense mutations and dnsSNPs in the new species was 3.1 and 2.9 times higher than those in the typical Malayan pangolins. The number of both missense mutations and dnsSNPs were in the sequence of MJ2 > MJ3 > MJ4 (**Fig. 5d–e** and **Additional File 1: Table S18**). Interestingly, both the typical Malayan pangolins and *Manis*_1 harbored much more homozygous LoFs than heterozygous ones. But this was opposite for the missense mutation and dnsSNPs. Additionally, homozygous missense mutation and dnsSNPs in *Manis*_1 were 10.8 and 9.5 times those in typical Malayan pangolins, although this was not found for the LoFs (**Additional File 1: Table S18**). Such difference was also observed for the ratio of mutations to nonsynonymous SNPs and mutational load (**Additional File 1: Figs. S19–20**). We found that the frequency of all three types of homozygous mutational loads in the ROH regions was higher than in the non-ROH regions (**Fig. 5f**).

## Discussion

### Large population differentiation in Malayan pangolins

The Malayan pangolins have been divided into two genetically distinct populations, the continental distributed MJA and MJB distributing on Southeast Asian islands, and each of the two populations consisted of two and three subpopulations, respectively^17^. This suggests differential evolution in a relatively stable continental environment and dynamic island environment. However, our large-scale analysis suggest that the Malayan pangolin consists of four genetically distinct clades, *Manis*_1, MJ2, MJ3 and MJ4. Specifically, the continental and island populations proposed in Hu et al. (2020a)^17^ were connected by addition of our samples, suggesting it to be a continent-peninsula-islands metapopulation corresponding to the clade MJ4 (**Figs. 1c–d** and **3c**). The clade MJ3 was made up of samples from the northwestern and southern Borneo and Java island despite isolation by sea (**Fig. 3b–c**). The clade MJ2 also showed great genetic differentiation from MJ3 (*F*_ST_ = 0.12) (**Figs. 1** and **3**), although it represented samples from the northeast part of the same large island (**Figs. 3b–c**). These divergence patterns suggest that terrestrial habitat seems more essential than the marine boundary to the genetic structure of Malayan pangolin.

Furthermore, the three clades of typical Malayan pangolin we have defined have different geographical locations (**Fig. 3**) and demographic trajectories (**Fig. 4e–f**) with no admixture (**Additional File 1: Fig. S5**). Genomic differentiation reached and even exceeded the subspecies level of the giant panda (*F*_ST_ = 0.14)^28, 36^ and leopards in Africa^37^(**Fig. 1h**), although subspecies differentiation can be even deeper in species like the tiger (*F*_ST_ = 0.32)^33^ and Chinese pangolin populations (*F*_ST_ = 0.49)^17, 38^, supporting the notion that the three clades of Malayan pangolin are evolutionary significant units and should be regarded as different conservation units (CU).

### Evidence for a new species

The hypothesis of a fifth Asian pangolin species was previously proposed according to phylogenetic analysis of partial mitochondrial genes^26, 39^. We have lent support to this hypothesis with our analysis presented here. The most compelling evidence was that the clade *Manis*_1 had the greatest *F*_ST_ when compared to the typical Malayan pangolin three clades, and their mitochondrial haplotypes were clustered apart from the other Malayan pangolin specimens and with those reported in Zhang et al. and Hu et al. ^26, 39^. We therefore propose that it is very likely that *Manis*_1 is not a clade of Malayan pangolin, but a new pangolin species.

Our analysis showed the following evidence for this proposal: First, *Manis*_1 formed a distinctive monophyletic clade in phylogenetic trees reconstructed using both the whole-genome SNPs and mitogenomes (**Fig. 1a**) lying between *M. crassicaudata* and the clade of *M. culionensis + M. javanica*. Second, the *F*_ST_ between *Manis*_1 and typical Malayan pangolin clades ranged from 0.71 to 0.75, reaching species-level when compared with recognized species in *Manis,* which was 0.79 between *M. pentadactyla* and *M. crassicaudata*, 0.82 between *M. pentadactyla* and *M. javanica*, and 0.74 between *M. crassicaudata* and *M. javanica* (**Fig. 1f**). Third, *Manis*_1 experienced a distinct demographic history compared to typical Malayan pangolin clades (**Fig. 4e–f**). Fourth, the skull of a sample of *Manis*_1 was consistent with Malayan pangolin according to morphological diagnostic characters (the ratio of NBL to GLS > 1/3)^22^. However, it markedly differed from typical Malayan pangolins at multiple positions, particularly the hamulus of pterygoid bone (**Fig. 2a**–**b**). On the CVA scatter plot of 75 landmarks it significantly separated apart from skull specimens of typical Malayan pangolins (**Fig. 2c**) and the four Asian pangolins in morphological space (**Fig. 2d**). The combined genetic and morphological evidence above supports that *Manis*_1 is a fifth species within the Malayan pangolin under both the phylogenetic species concept^40^ and the gen-morph species concept^41^.

We recommend that the phylogenetic relationships in Asian *Manis* be revised as: *M. pentadactyla +* (*M. crassicaudata +* (*Manis_*1 + (*M. culionensis + M. javanica*))). Inference of demographic history of Malayan pangolin using MSMC2 often sets generation time (g) to 1 or 7 years^17, 27^. However, g of the species should be > 2 years but may not exceed 5 years according to its reproductive parameters in captivity^42, 43^. We used 3 and 5 years as the two boundaries and showed the divergence time between *Manis*_1 and typical Malayan pangolin was between 3 and 2 Mya, earlier than the divergence between *M. javanica* and *M. culionensis* (1.7 Mya [2.5–0.4]^24^).

One sample (P442988) showed nuclear-mitochondrial discordance. Its mitochondrial haplotype belonged to the HapGroup III as *Manis*_1, but the nuclear genome completely belonged to the typical Malayan pangolin (**Figs. 1a** and **3c**). It is very likely to have resulted from historical introgression of hybrids with maternal *Manis*_1 repeatedly backcrossed to paternal typical Malayan pangolin ^44–48^. This sample originated in northeastern Borneo. We suspect that the new species might have been distributed in northeastern Borneo or its adjacent areas.

The sample PA belonging to *Manis*_1 (**Figs. 1a** and **3c**) was previously identified as *M. culionensis* by Cao et al. (2021)^49^. However, according to the ML tree constructed with complete mitogenomes^49^, it does not form a monophyletic clade with *M. culionensis*, but is actually outside the *M. culionensis* and *M. javanica* clade. This is concordant with the phylogenetic relationships between *Manis*_1 and the two species reconstructed in our study (**Fig. 1a**), suggesting it more likely a member of *Manis*_1 instead of *M. culionensis*.

### Genetic risk and conservation implications

Genetic diversity is an important factor for understanding target species^50, 51^ and informing conservation initiatives. Pangolins in southeast Asia has been heavily impacted by poaching and illegal trade^19, 25^. We show that pangolins inhabit almost the entire region with strong habitat heterogeneity and show higher levels of diversification that previously reported. Based on genome-wide and mitochondrial genome data from a large sample representing genetically distinctive clades, we assessed genetic diversity for the typical Malayan pangolin and the new species *Manis*_1. The genome-wide *He* varied greatly in typical Malayan pangolins, but the majority was at a moderate level (**Fig. 5a** and **Additional File 1: Table S5**), suggesting a substantial basis for sustainability. The clade MJ4 shows the most genetic variation among the typical Malayan pangolins is likely due to its distribution across a variety of habitats from continent, the peninsula, and islands, leading to corresponding spatial genetic structure (**Fig. 1c–d**). However, it is noteworthy that the mainland populations have the least genomic heterozygosity and highest inbreeding level. These populations require additional attention to safeguard genetic diversity. For *Manis*_1, the average *He* (0.00132) was considerably lower than those of the typical Malayan pangolins (MJ2–MJ4, 0.00180) and the Chinese pangolin (0.00182)^38^. This level is moderate among selected endangered species, but indicates risk of collapse given their population size and distribution range.

Inbreeding is an important contributor to the loss of genetic diversity, recent inbreeding often leads to long ROH on the genome, and ancient inbreeding leads to short ROH^52^. We observed that the majority of ROH in all clades of typical Malayan pangolin and *Manis*_1 was medium-sized (100 kb to 1 Mb), suggesting the pangolins had experienced high levels of ancient inbreeding instead of recent inbreeding in the population history (**Fig. 5b**). It tends to rule out that recent anthropogenic disturbance has made severe impacts on genetic diversity, indicating that pangolins could rebound to a sustainable status by eliminating impact factors such as poaching, illegal trade, and habitat destruction.

Accumulation of deleterious mutations is a key genetic factor that can be used for evaluating the decrease of fitness small populations^53, 54^. We found that the quantity of LoF in *Manis*_1 was comparable to the typical Malayan pangolin. However, *Manis*_1 had greater proportion of homozygous loads (**Additional File 1: Fig. S20**). This suggests *Manis*_1 might potentially have a higher fitness cost^33^, and should receive special attention.

## Conclusions

The Malayan pangolin of southeast Asia has diverged into three distinct clades. MJ2 inhabits northeast Borneo, MJ3 inhabits northwest to south Borneo and south Sumatra, and MJ4 from the south part of Asia continent to Sumatra Island via Malayan peninsula. MJ4 differentiated into at least 17 subpopulations with gene flow between neighbors. *Manis*_1 is a new species that is more closely related to the Philippine pangolin than the typical Malayan pangolin, but a more distant relative of the Indian pangolin. The phylogenetic relationship of Asian pangolins should be revised as *M. pentadactyla +* (*M. crassicaudata +* (*Manis_1* + (*M. culionensis + M. javanica*))). Genetic diversity of all clades of typical Malayan pangolin and *Manis*_1 was moderate compared to other endangered species, with the exception of mainland subclade in MJ4. Recent inbreeding is relatively low, suggesting anthropogenic factors did significantly weaken the basis for sustainability. However, eliminating impacts of poaching, illegal trade and habitat destruction, may enable pangolins to rebound to sustainable status. Finally, ecological study and genetic and morphological characterization of the new species and evolutionarily significant subpopulations of typical Malayan pangolins should be prioritized to accurately support conservation, avoiding misidentification of forensic samples and mistaken interbreeding of rescued animals for rehabilitation. Special attention should be given to mainland Malayan pangolins due to their relatively low genetic diversity, high inbreeding, and fitness cost.

## Methods

### Sample collection and Ethics Statement

A total of 594 pangolin individuals were collected and sequenced, including 14 individuals collected from natural history museums in the USA via the University of Washington Center for Environmental Forensic Science, 131 individuals collected from College of Wildlife and Protected Area in Northeast Forestry University, Heilongjiang Province, China and 449 individuals from the Guangzhou Wildlife Rescue Center, Guangdong Province, China. Samples were collected under the supervision of Institutional Review Board of BGI and permission was obtained when necessary (BGI-IRB: E21056-T1). Whole-genome sequencing data of 73 Malayan pangolin individuals^17, 27^ were downloaded from National Center for Biotechnology Information (NCBI, **Additional File 1: Tables S1** and **S4**) for downstream population genetic analysis in this study. In addition, 19 mitochondrial genome sequences from eight living species of Manidae (*Phataginus tricuspis*, *Phataginus tetradactyla*, *Smutsia gigantea*, *Smutsia temminckii*, *Manis pentadactyla*, *Manis crassicaudata*, *Manis culionensis* and *Manis javanica*), and the outgroup cat (*Felis catus*)^55^and dog (*Canis lupus familiaris*)^56^ were downloaded from NCBI for species identification and mitochondrial phylogeny construction (**Additional File 1: Table S15**). Sixty-two *COX1* gene sequences and 62 *Cytb* gene sequences were also downloaded from NCBI for haplotype network construction^17, 24^(**Additional File 1: Table S17**). Except for Malayan pangolins, three Tree pangolins, two Chinese pangolins and one Giant pangolin were identified by mitochondrial genes.

### Morphometry analysis

To test morphometric variation of 42 skulls of the sequenced pangolins. Ten morphological indexes were measured with standard methods^35^, namely the greatest length of skull (GLS), basal length (BL), palatal length (PL), the greatest cranial breadth (GCB), interorbital breadth (IOB), length of auditory bulla (LAB), high of brain case (HB), nasal length (NBL), greatest breadth of nasal (GBN) and mandibular length (ML) (**Additional File 1: Table** S7). The value of each index was divided by the GLS, and a Principal Component Analyses (PCA) was performed using the FactoMinR package with the RStudio software (version: 1.1.463).

We also scanned these skulls using Micro-CT (Scanco medical, vivaCT 80, Switzerland). A total of 75 three-dimensional landmarks^57^ were located on pangolin skulls using Stratovan CheckPoint^®^ software. The similar data of 32 skulls from four Asian pangolins^57^ were included in following analysis (**Additional File 2: Extended Data Table 1**). The data of landmarks were imported in MorphoJ^58^ to analyze morphological differences of skulls. We performed generalized Procrustes analysis (GPA)^59^ to eliminate effects of orientation, position and size, and got centroid size (CS) values^60^ and matrices of shape coordinates (Procrustes coordinates)^61, 62^. The shape features of each group were distinguished by Canonical variable analysis (CVA). The distance of each group from the sample mean was measured in Mahalanobis distance. Procrustes distances were used to quantify the distance of shape differences between groups, which was an absolute measure of the magnitude of shape differences between groups.

### Library preparation and sequencing

For each pangolin, genomic DNA was extracted from muscle and integumentary appendages samples using a standard phenol/chloroform extraction^63^. DNAs with sufficient concentration (> 500 μg) and high fragment integrity (agarose gel electrophoresis) were selected for library construction. Paired-end sequencing libraries with an insert size of 350 bp were constructed according to the manufacturer’s instructions for sequencing on the DNBSEQ T1 platform (BGI, Shenzhen, China).

### Species identification and mitochondrial Phylogeny Construction

For species identification, we generated the whole mitochondrial genome sequence for each individual with NOVOPlasty^63^(version: 4.3.1). We concatenated 13 mitochondrial coding genes (*ND1*, *ND2*, *ND3*, *ND4*, *ND 4L*, *ND5*, *ND6*, *ATP6*, *ATP8*, *COX1*, *COX2*, *COX3*, *Cytb*) to form a super gene with 668 pangolins, one cat, one dog and eight species of Manidae (*N* = 687). We aligned the dataset of super gene sequences with MEGA^64^(version: X). The alignment was partitioned using PartitionFinder^65^(version: 2.1.1). The analysis recommended splitting the dataset into one partition and proposed the best-fit nucleotide substitution model ‘GTR+I+G’ for the phylogenetic analysis. We then estimated a maximum-likelihood (ML) tree with 1000 bootstraps using IQ-TREE^66^(version: 1.6.6) and a Bayesian inference (BI) using MrBayes^67^(version: 3.2.5). Tree layout was generated using the online tool Interactive Tree of Life (iTOL, http://itol.embl.de).

For revealing population identity, we generated a haplotype network using the Median-Joining method in PopART^68^(http://popart.otago.ac.nz/). Haplotype network (Network_717) was construct with joint *COX1*-*Cytb* gene of 658 individuals in this study and 59 previously available samples with known origins, including Thailand, Borneo, Java, Singapore/Sumatra, Indonesia, and Malaysia. Given that two new suspected pangolin haplotypes (H04: KT428139.1, H08: KT428143.1) with short mitochondrial *COX1* (600 bp) and *Cytb* (399 bp) gene sequences were found in Hu et al.’s study^26^, we further generated another haplotype network (Network_660) for joint sequence (999 bp) of these two pangolins and the other 658 individuals by the above method.

### Read mapping and variants calling

Raw sequence reads were aligned to the *M. javanica* reference genome (YNU_ManJav_2.0, Genbank: GCA_014570535.1) using the Burrows-Wheeler Aligner^69^(BWA, version: 0.7.10-r789) with default parameters. Next, Picard tools (http://picard.sourceforge.net) (version: 2.1.1) and Genome Analysis Toolkit^70^(GATK, version: 4.0.3.0) were employed to process and filter the BAM format alignment file, including alignment position sorting, duplicated reads marking, local realignment, and base quality recalibration. Individuals sequencing coverage was calculated with Samtools^71^(version: 1.3). Individuals with coverage < 80% were discarded. Raw variant calling for each individual was performed generating a Genome Variant Call Format (gVCF) file using GATK with HaplotypeCaller. Lastly, joint calling was then performed to combine all gVCF files into the raw population Variant Call Format (VCF) file.

Hard filtering was performed on the raw SNP variant set with parameters of “QUAL < 30.0 || QD < 2.0 || FS > 60.0 || MQ < 40.0 || MQRankSum < -12.5 || ReadPosRankSum < -8.0”. In addition, we removed SNPs in scaffolds smaller than 100 kb (2.02% of the reference genome). SNP sites with extreme depths value (> 99.75% and < 0.25%) were then filtered. Subsequently, loci with a quality value (PL) greater than 20 and a missing rate less than 0.1 were retained using VCFtools^72^(version: 0.1.13). The final high-quality SNP set was used for further population genetic analyses. For PCA, admixture, and phylogenetic analysis the SNP set was pruned with VCFtools and the parameter “*thin 2000*” which retains one SNP every 2000 base pairs. The unthinned SNP dataset was used for the remaining analyses including *He,* inbreeding, mutational loads, etc. We inferred the family relationship among all Malayan pangolins with KING^73^(version: 2.2.7) to remove the potential consanguineous individuals. We retained one of close pairs (parent-offspring, monozygotic twin, full-sibling and 2nd Degree) and all unrelated individuals for the subsequent analyses (total 598 individuals).

All variants were annotated using the perl script “*annotate_variation.pl*” of ANNOVAR^74^(version: 2015-12-14), including exotic, nonsynonymous, synonymous, UTR, intronic, intergenic, splicing, and non-coding RNA (ncRNA).

### PCA, Phylogeny, and Admixture Clustering Analyses

PCA was performed on the pruned SNP set using Genome-wide Complex Trait Analysis^75^(GCTA, version: 1.91.4beta3). Due to the large genetic distance between *Manis*_1 individuals (MJ-DCW-89/MJ-DCW-116) and the other 596 individuals, we further performed PCA analysis on the Malayan population except for these two samples. A ML phylogenetic tree was constructed with 1000 bootstraps using IQ-TREE and the tree layout was generated using the online tool iTOL. The population structure was analyzed with the cluster number K ranging from 2–20 by ADMIXTURE^76^(version: 1.3.0).

### Population differentiation and gene flow

We performed the f3-test using qp3Pop implemented in ADMIXTOOLS^77^(version: 5.1) to determine if one population was an admixed population of the other two. A Z-score with the absolute value greater than 3 was used to determine the significance. We then used Weir and Cockerham’s *F*_ST_ to estimate the population differentiation. All bi-allelic SNPs were used for the calculation of genome-wide *F*_ST_ between each pair of the populations using the VCFtools “*--fst-window-size 200000 --fst-window-step 100000*” tool.

We applied the classic ABBA-BABA test (D statistics) to indicate the occurrence of gene flow by the software POPSTATS^78^ with the command “-*informative*”. The four populations and the outgroup were set to be ((A, B), (X, Y)), and the A, B and X were among each of *Manis*_1, MJ2, MJ3 and MJ4. Chinese pangolin was used as the outgroup (Y) according to the phylogenetic tree. A significant deviation of D-statistics from zero (|Z-score| > 3) was due to different numbers of shared sites between X and either A or B.

Identification of sharing of identity by descent (IBD) between individuals in each pair of populations could improve the accuracy of admixture inference ^79^. IBD was calculated by the RefinedIBD (version: 16May19. ad5) with default parameters, except the length of IBD fragment (*length = 0.1*). We compared the total length of IBD fragments (IBD > 1 Mb and 100 kb < IBD < 1 Mb) and their percentage on the reference genome among populations to evaluate the degree of gene flow.

### Demographic analysis

We first used MSMC2^80^(version: 2.1.2) to infer the separation among four populations. Genotype phasing was performed by using the Beagle (version: 5.0) software with default parameters before the MSMC2 inference. MSMC2 was performed for six independent replications with two randomly selected samples from each population. Parameters for MSMC2 calculations were as follows: *--skipAmbiguous -I 0-4, 0-5, 0-6, 0-7, 1-4, 1-5, 1-6, 1-7, 2-4, 2-5, 2-6, 2-7, 3-4, 3-5, 3-6, 3-7 -i 20 -t 6 -p ‘10*1+15*2’*. The mutation rate and generation interval of the *M. javanica* we used here was 1.47×10^-8^ per site per generation^17^ and 3–5years^27^.

### Genetic diversity and Runs of Homozygosity (ROH)

Whole-genome genetic diversity, including heterozygosity (*He*), and nucleotide diversity (π) in a nonoverlap 50 kb window were computed with VCFtools. ROHs were identified for each individual using the “*run of homozygosity*” function in PLINK^81^(version: 1.90b4.6). We ran sliding windows of 20 SNPs on the VCF files of each scaffold, requiring at least one SNP per 50 kb for ROH fragments more than 10 kb in length. In addition, we allowed for the maximum of one heterozygous SNP per window. We compared the total length of ROH fragments (ROH > 1 Mb and 100 kb < ROH < 1 Mb) and their fraction in the reference genome (F_ROH_) among populations.

### Mutational load

We identified derived mutational loads with *F. catus* as the reference genome. A deleterious mutation is an amino acid change in a protein that was predicted to be harmful to its function. For Loss-of-Function (LoF) mutational variants, we selected derived mutations in coding regions of each pangolin individual via annotation by SnpEff^82^(version: 4.3). LoF variants included *splice_donor_variant*, *splice_acceptor_variant* and *stop_gained*. For missense variants, we identified by the type ‘*missence_variant*’ annotated through SnpEff. For deleterious nonsynonymous SNPs (dnsSNPs), we used the of parameter “*--aamatrixfile grantham matrix*” in the package ANNOVAR to obtain Grantham Score (GS) for nonsynonymous variants. When GS L≥L150, the mutation can be designated as deleterious^83, 84^. We counted the number of mutational variants and the ratio of mutational variants to total nsSNPs to compare the accumulation mutational variants of each population. The ratio of homozygous (two per site) to (homozygous (two per site) plus heterozygous sites (one per site)) for mutational variants was calculated to estimate the level of mutational load. We further compared the ratio of mutational variants in the ROH and non-ROH genome regions in each individual to detect a purging effect.

## Acknowledgements

This study was jointly supported by the Fundamental Research Funds for the Central Universities (No. 2572020DR10), National Key Program of Research and Development, Ministry of Science and Technology (No. 2022YFF1301500), Hunan Provincial Forestry Science and Technology Innovation Plan Project (No. XLK201915), Rare and Endangered Species Investigation and Industry Regulation Project (No. 2020070209), Guangdong Provincial Key Laboratory of Genome Read and Write (No. 2017B030301011), and funding for generation of the sequenced reads for individuals of known geographic origin was supported by the USAID Wildlife Crime Tech Challenge to HJK and SW. We thank George Amato and Eleanor Hoeger at American Museum of Natural History (AMNH), Nicole Edmison at Smithsonian Institute, Louise Tomsett at Natural History Museum (NHM) London, and Sharlene Santana, Sharon Birks, and Jeffrey Bradley at Burke Museum for their assistance in acquiring the specimens of known origin. Thanks to Professor Shibao Wu for his guidance and help in morphology. Finally, we are thankful to the China National GeneBank for producing the sequencing data.

## Author contributions

Y.X., B.L., H.L., T.L., Y.H. and L.L. designed the research. Y.H., L.L., Z.D., H.G., Y.Y., Z.W., W.Z., S.B. and J.Y.L. provided the pangolin samples. Y.M., X.Z., K.W., B.Y.L., L.C., C.W. and J.H.L. led and finished the genomic DNA preparation. H.M.L., S.Q.W., S.Y., J.G.W., X.W., H.R.L., J.F., W.L., X.N., J.Y., L.H., DY.Y., J.C., Y.X.W., X.C. and Y.T.H. led and finished the genome sequencing. H.J.K. sequenced samples with known geographic locations. Y.X., C.G., B.L., H.X., Y.Y.W., Y.M.H., J.L., S.Y.L., Y.J., S.S.L. and J.Y.W. collected skull data. Y.X., T.L., Q.W., M.S., C.G., Z.Q. and B.L. analyzed and interpreted the data. Y.X., B.L., T.L., Q.W., M.S. and C.G. wrote the manuscript. Y.X., B.L., T.L., Y.H., S.L.L, K.F. and S.K.W. review and edited the manuscript.

## Data availability statement

The data that support the findings of this study have been deposited into CNGB Sequence Archive (CNSA)^85^ of the China National GeneBank DataBase (CNGBdb)^86^ with accession number CNP0002036.

## Supplementary Information

Additional file 1: Supplemental figures 1–20 and Supplemental tables 1–18.

Additional file 2: Extended Data Table 1: Raw landmark coordinates.

